# M-CAMP™: A cloud-based web platform with a novel approach for species-level classification of 16S rRNA microbiome sequences

**DOI:** 10.1101/2021.08.25.456838

**Authors:** Andrew E. Schriefer, Brajendra Kumar, Avihai Zolti, R Preetam, Adam Didier, MG Nirmal, Nofar Nadiv, Michael Perez, Santosh Kumar Mahankuda, Pankaj Kumar, G T Greeshma, Aaron Tenney, Maureen Bourner, Shira Lezer, Fei Zhong, Michal Daniely, Yang Liu

**Author notes:** To Whom Correspondence should be directed. Joint first authors (Andrew E. Schriefer, Brajendra Kumar).

## Abstract

The M-CAMP™ (Microbiome Computational Analysis for Multiomic Profiling) Cloud Platform was designed to provide users with an easy-to-use web interface to access best in class microbiome analysis tools. This interface allows bench scientists to conduct bioinformatic analysis on their samples and then download publication-ready graphics and reports. The core pipeline of the platform is the 16S-seq taxonomic classification algorithm which provides species-level classification of Illumina 16s sequencing. This algorithm uses a novel approach combining alignment and kmer based taxonomic classification methodologies to produce a highly accurate and comprehensive profile. Additionally, a comprehensive proprietary database combining reference sequences from multiple sources was curated and contains 18056 unique V3-V4 sequences covering 11527 species. The M-CAMP™ 16S taxonomic classification algorithm was validated on 52 sequencing samples from both public and in-house standard sample mixtures with known fractions. Compared to current popular public classification algorithms, our classification algorithm provides the most accurate species-level classification of 16S rRNA sequencing data.

## Introduction

Over the past decade, with the advancement of sequencing technologies and metagenomics, the science of studying microorganisms in certain environmental habitats is becoming increasingly important. Modern metagenomic studies allow researchers to uncover numerous previously unknown species from an environmental niche, or to classify large amounts of DNA sequences to known species. It has been compared to ‘a reinvention of the microscope’, allowing scientists to create a spectrum of microbials without isolating individual microbes (Council, 2007). A wide variety of topics have been studied, with environmental focuses such as seawater (Venter et al., 2004) to acidic mine drainage (Tyson et al., 2004) to the microbiome’s role in diseases in the human body (Lloyd-Price et al., 2019) (Zhou et al., 2019) (Fettweis et al., 2019).

16S ribosomal RNA sequencing is one of the primary means of classifying bacterial species (Woese, 1987). 16S-seq is a cost effective and high-throughput way of characterizing microbiome samples. Many algorithms have been developed to facilitate 16S rRNA classification (Gao et al., 2021), such as Quantitative Insights into Microbial Ecology (QIIME) (Caporaso et al., 2010), Kraken (Wood & Salzberg, 2014), MAPseq (Matias Rodrigues, Schmidt, Tackmann, & von Mering, 2017), Spingo (Allard, Ryan, Jeffery, & Claesson, 2015), IDTAXA (Murali, Bhargava, & Wright, 2018), etc. These algorithms follow two distinct approaches. The first approach is to use an alignment method to reference 16S sequences. This approach gives very good accuracy down to the species level with a higher cost of computational resources and greater difficulty in identifying new bacteria. Alignment based tools include MAPseq and the vsearch classifier in QIIME (Rognes, Flouri, Nichols, Quince, & Mahé, 2016). The second approach is to use kmer-based classification, often coupled with a machine-learning algorithm using short kmers as features. These approaches are less computationally intensive and can generalize more easily to bacterial sequences not contained in the reference database, but with reduced accuracy at the species level. Kmer based tools include Kraken, Spingo, IDTAXA and sklearn classifier in QIIME. Since 16s sequencing has much smaller file output than WGS output (typically 10-100 times smaller), the additional computation cost is not as significant; however, the increased accuracy of species level organism identification is much more beneficial for scientists. For example, it is well known that species-level classification with 16S rRNA gene sequencing of *Salmonella enterica* is challenging and is critical to researchers (Grim et al., 2017).

Another key factor in the 16S classification is the 16S sequence reference database. There are several well-known databases, including Greengenes (DeSantis et al., 2006), SILVA (Quast et al., 2013), RDP (Cole et al., 2014), along with 16S sequences deposited in NCBI. Despite these high-quality resources, there remains need for further manual curation to achieve accurate species-level classification (Srinivasan, Karaoz et al. 2015).

In this paper, we present a novel approach combining alignment and kmer classification to better characterize 16s rRNA sequencing data. We fine-tuned the alignment algorithm to balance computation time and alignment results using standard mixtures to achieve optimal classification results. Additionally, we manually curated a 16S rRNA database containing the most up-to-date references from multiple sources. With the M-CAMP™ Platform, we are making this tool available to the scientific community to facilitate microbiome research. The web-based platform allows user to upload their sequencing data and perform taxonomic classification and comparative analysis.

## Methods

### AMP™ Cloud Platform

The M-CAMP™ Cloud Platform is a cloud-based web application for classifying species-level bacterial and archaeal taxonomic profiles from 16S-seq data. The platform, which is provided as a web-service for the scientific community, delivers best-in-class taxonomic profiling performance by combining two components in the classification algorithm. The first is a novel algorithm which generates precise species-level classifications for known 16S sequences, offering better coverage for ambiguous or unknown 16S sequences than were previously available. The second is a curated database, which resolved ambiguities in publicly available data and has been specifically designed for the data generated by the Sigma-Aldrich microbiome laboratory. The web platform is hosted on Google Cloud Platform and uses Java EE technology. Users can easily access and submit their sample data and perform analyses from microbiome projects.

### Classification Algorithm

Our classification approach consists of two steps, a preprocessing step removing low quality sequences and a classification step accurately assigning sequences to proper species. Input fastq reads are preprocessed using the following steps below prior to taxonomic classification using commands from BBTools (Bushnell, Rood, & Singer, 2017). The preprocessing steps are performed to reduce false positive taxonomic classifications. The first step is quality trimming using the default parameters of bbduk.sh. The second step is trimming primer sequences from the 5’ end of the forward and reverse reads by using reformat.sh to trim a number of bases equal to the length of the targeted 16S primers used to generate the sequencing sample. Primer sequences are removed because PCR errors during library preparation of Illumina amplicon sequencing can introduce errors into the primer regions of the read (Zhang et al., 2019). The third step is merging the paired reads by overlap using default parameters of bbmerge.sh. If the input fastq files are single-ended the pair merging step is skipped. All reads that fail to merge by overlap are discarded from any further processing and they are reported in the output as “unclassified”. The final step is removing chimeric reads using vsearch (Rognes et al., 2016) with the –unchime_ref command.

At the classification step, all reads passing the preprocessing stage are aligned to the primer-specific 16S gene sequence. The output bam file is then processed with a custom script which classifies each read based on the lowest common ancestor of its multimappings. Reads with no mappings are then classified using Kraken2 which uses a kmer-based approach to classifying reads (Wood, Lu, & Langmead, 2019). Only the genus level classifications are included from the Kraken2 results to reduce over-classification error by kraken on sequences that are not present in the reference database. Classifications from Kraken and the LCA classifications from sequence alignment are combined into a single report and any taxonomic labels with relative abundance less than 0.01% are removed.

### Classification Database

The 16S rRNA database used by our classification algorithm involves a two-step process. The database build process takes as input full length 16S gene sequences from different reference sets including Refseq prokaryote genomes, Refseq Targeted Loci 16S ribosomal RNA project (O’Leary et al., 2016), and SILVA release 138. The process begins by running the input full-length 16S sequences through ecoPCR from obitools (Boyer et al., 2016) using the V3-V4 primer sequences Bakt_341F (CCTACGGGNGGCWGCAG) and Bakt_805R (GACTACHVGGGTATCTAATCC) (Herlemann et al., 2011). ecoPCR is run, allowing up to two mismatches and disallowing mismatches on the last two bases of the primers. The obiuniq command is used to combine identical predicted amplicon sequences into a single reference sequence. Each reference sequence then has a list of NCBI taxids for all the predicted amplicon sequences with identical sequences. The reference sequence taxonomy label is creating using an α-majority LCA algorithm on these NBCI taxid counts as described previously (Hanson, Konwar, & Hallam, 2016) with α=0.8.

Second, the database combination process involves two databases: a base and an addition. The purpose of the addition database is to add sequences to the base to create a more comprehensive database. Sequences from the addition database are aligned to the base database using vsearch. Any sequence in the addition which maps to the base at sequence identity greater than 0.99 are removed and the rest of the sequences are added in. This combined database becomes the base and the next database to be added in becomes the addition. This process continues until all the databases have been merged.

The first base database used to generate the M-CAMP™ V3-V4 combined database is generated from the RefSeq prokaryote genomes. DNA sequences of 16S genes were extracted from the genomes using barrnap (Seemann, 2013) and the sequences are annotated with the taxids of the source genome. The RefSeq Targeted Loci project is added next and SILVA release 138 is added last. The final merged database contains 18056 unique V3-V4 sequences covering 11527 species 3359 genera. Compared to the SILVA 138 database processed with the V3V4 primers which has 185418 of sequences covering 48113 species and 4723 genera. The number of sequences in the full SILVA database is larger but many the sequences have questionable taxonomic classifications which may cause issues on classification at species level. Our merging process ensures that only the highest quality taxonomic information is used for classification.

### Comparison Algorithms

The following 16s-seq classification algorithms were downloaded and installed for comparison to our classification method. When possible, the software was installed using bioconda (Gruning et al., 2018). When that was not possible, the installation instructions provided by the software maintainers were instead followed. The comparisons were run on a CentOS Linux server with 16 cores and 64 GB of memory.

1. Qiime2 version 2019.10 (Bolyen et al., 2019) using SILVA 138
2. Spingo version 1.3 (Allard et al., 2015) using RDP 11.2
3. Mapseq version 1.2.3 (Matias Rodrigues et al., 2017) using Mapref 2.2b
4. Idtaxa version 2.16 (Murali et al., 2018) using SILVA 138
5. Kraken2 version 2.0.8 (Wood et al., 2019) using SILVA 138

There are a handful of database sources used by these algorithms. These include SILVA (Quast et al., 2013), RDP (Cole et al., 2014), Greengenes (DeSantis et al., 2006), and RefSeq TargedLoci. For the benchmarking, a variety of database sources were used to run the various algorithms. In general, the author-suggested or default database was used for the benchmark process. The Greengenes database was avoided because of its lack of recent updates. The choice to not explicitly standardize the database for each algorithm was made because most users will follow defaults;the default represents the real-life performance most users will experience. Many of the algorithms rely on specially formatted database files which are difficult to reproduce and must be provided by the author. In these cases, there is no choice in which database to use. For example, the SPINGO github only provides one database which is a derivative of RDP. Detailed commands used to run each classification method are available in the **Supplement**.

### Validation Samples

To evaluate performance of classification algorithms, including ours and others published previously, we used a combination of samples from public resources and in-house samples from our Sigma-Aldrich R&D team. These samples included the Sigma-Aldrich Microbial community DNA mix (Sigma-Aldrich, Catalog: MBD0026), synthetic mixtures from other commercial vendors, single bacterial standards (Sigma-Aldrich, Catalog: MBD0001 – MBD0024) samples, and double bacteria mixture samples. All the samples used in this study were sequenced by the Sigma-Aldrich R&D team. The samples were amplified using the V3V4 primers and sequenced on an Illumina MiSeq.

Previously published 16S Illumina sequencing data of microbial mixtures were also included. Schirmer MUB and MB (Schirmer et al., 2015) mixtures contain 59 bacteria and archaea strains sequenced using different Illumina library preparation methods. The MB samples contained mixtures of the microorganisms in even uneven proportions and the MUB samples contain mixtures of uneven proportions. Of those all the samples sequenced in the Schrimer manuscript we selected 10 samples using both the V3V4 and V4 amplicon. Tourlousse Ec and Q samples are mixtures of 15 bacterial genome sequenced using the V3V4 amplicon (Tourlousse et al., 2017). This mixture was designed to be used as a synthetic spike-in to 16s samples for purposes of relative abundance calculation and quality control. We include these samples in our validation set because they contain microbial species not found in other synthetic communities. Finally, we tested data from Gohl M. et al, consisting of several sequencing samples of the Human Microbiome Project Genomic Mock Community B (Gohl et al., 2016) (HM-276D, Even, High Concentration, v5.1H, and HM-277D, Staggered, High Concentration, v5.2H). These samples were sequenced using the V4 amplicon and they consisted of the same 20 bacteria present in the ATCC mixtures.

We sequenced 10 human stool samples acquired from Lee Biosolutions (Catalog 991-18). These samples contain complex communities from real human donors that are more representative of real research samples than the synthetic communities included in our validation samples. The purpose of sequencing these stool samples is to assess how well each method generalizes to complex real-world samples. The bacteria present and their relative abundances are completely unknown for these samples.

A detailed description of all the groups of microbiome samples used in the evaluation are available in **Table 1**. Information about each sequencing sample used is available in **Supplemental Table 1**

**Table 1:** Information on Validation Sequencing Sample Groups.

| Mixture | # Microbes | Is Even Mixtures | # Samples | Source | Sequencing Data Source |
| --- | --- | --- | --- | --- | --- |
| Zymo | 8 | Yes | 9 | ZymoBIOMICS | MilliporeSigma |
| MSA-1001 | 10 | No | 6 | ATCC | MilliporeSigma |
| MSA-1000 | 10 | Yes | 6 | ATCC | MilliporeSigma |
| Sigma-Aldrich Mix | 10 | No | 3 | MilliporeSigma Catalog MBD0026 | MilliporeSigma |
| Schirmer MB | 59 | Yes | 6 | https://doi.org/10.1093/nar/gku1341 | http://www.ebi.ac.uk/ena/data/view/PRJEB6244 |
| Schirmer MUB | 59 | No | 5 | https://doi.org/10.1093/nar/gku1341 | http://www.ebi.ac.uk/ena/data/view/PRJEB6244 |
| Tourloussie Ec | 15 | Yes | 1 | https://doi.org/10.1093/nar/gkw984 | https://www.ncbi.nlm.nih.gov/biosample/SAMN05200489/ |
| Tourloussie Q | 15 | No | 1 | https://doi.org/10.1093/nar/gkw984 | https://www.ncbi.nlm.nih.gov/biosample/SAMN05200490 |
| HM-276D | 20 | Yes | 2 | https://doi.org/10.1038/nbt.3601 | https://www.ncbi.nlm.nih.gov/biosample/SAMN04325243/ |
| HM-277D | 20 | No | 2 | https://doi.org/10.1038/nbt.3601 | https://www.ncbi.nlm.nih.gov/biosample/SAMN04325244 |
| Single Mixtures | 1 | Yes | 9 | MilliporeSigma MBD0001-MBD0024 | MilliporeSigma |
| Double Mixtures | 2 | Yes | 2 | MilliporeSigma MBD0001-MBD0024 | MilliporeSigma |

### Sequencing and Sample Prep Methods

DNA was extracted from Microbial Community Standard mixtures by MagMAX™ Total Nucleic Acid Isolation Kit (AM1840) on a KingFisher™ Flex Purification System platform (ThermoFisher). DNA libraries from all bacterial mixture samples were prepared in a dual amplification procedure; First a PCR amplification (25 cycles) was performed with the above mentioned V3V4 primers, followed by 8 additional PCR cycles with Nextera XT Index Kit v2 Sets A and/or D (Catalog #FC-131-2001 and FC-131-2004, Illumina). Libraries were pooled and sequenced using MiSeq V2 reagent kit, with Nano configurations, on an Illumina MiSeq sequencer, yielding paired-end 2 × 250 base reads. Wet lab preparation of the samples followed best-practice recommendations described here (Nearing, Comeau, & Langille, 2021).

### Classification Accuracy

We ran all 52 validation samples using our algorithm, and other comparison algorithms as described above. The species-level results were compared using the F1 score, precision, and recall. Precision is inversely related to the number of false positive bacteria and recall is inversely related to the number of undetected true positive bacteria. The F1 score is a combination of precision and recall, giving an overall measure of how well the calculated and the known taxonomic profiles match.

The F1 score, precision and recall were calculated using the python scikit-learn metrics module. Code to calculate the L2 (Euclidean) distance was adapted from another metagenomic methods benchmarking paper (Ye, Siddle, Park, & Sabeti, 2019). For each method, the output files for all samples were parsed into a data table containing relative abundances and NCBI taxonomy IDs. Any NCBI taxonomy ID present at less than 0.01% relative abundance was filtered from the results for purposes of calculating the evaluation metrics. When an algorithm was run using a different taxonomy than NCBI, such as Silva, the taxonomy names were mapped to NCBI taxonomy ids using the ete3 library (Huerta-Cepas, Serra, & Bork, 2016). Metrics were calculated for different ranks by summing up all the abundances of all descendant NCBI taxonomy IDs.

## Results

### Evaluation of Classification Accuracy

Here we summarize the results of all classification algorithms on the entire set of 52 known mixture samples. We ran every classifier algorithm on each sample and compared the predicted taxonomic profile with its known taxonomic profiles and relative abundances. We then assessed the performance by calculating the precision, recall, F1-score, Bray-Curtis distance, and L2 distance (otherwise known as the Euclidean distance) for each of the seven main ranks of the NCBI taxonomy for prokaryotes. The average values of the evaluation metrics for each method averaged across all the validation samples at multiple taxonomic levels are shown in **Supplement Table S3**. Boxplots containing all data points at the species, genus, and family levels for the M-CAMP™ pipeline and the comparison methods are shown in **Figure 2, Figure 3**, and **Figure 4. Figure 2** contains the F1 values which are calculated from the binary presence/absence of correct and incorrect microbes. **Figure 3** contains the L2-norm metric which is calculated using the relative abundance values of correct and incorrect microbes. **Figure 4** contains the Bray-Curtis distance metric, which is a popular measure of beta diversity. This distance was calculated using the relative abundance values of correct and incorrect microbes. These boxplots show that most methods perform about the same at the genus and family level.

**Figure 1:**
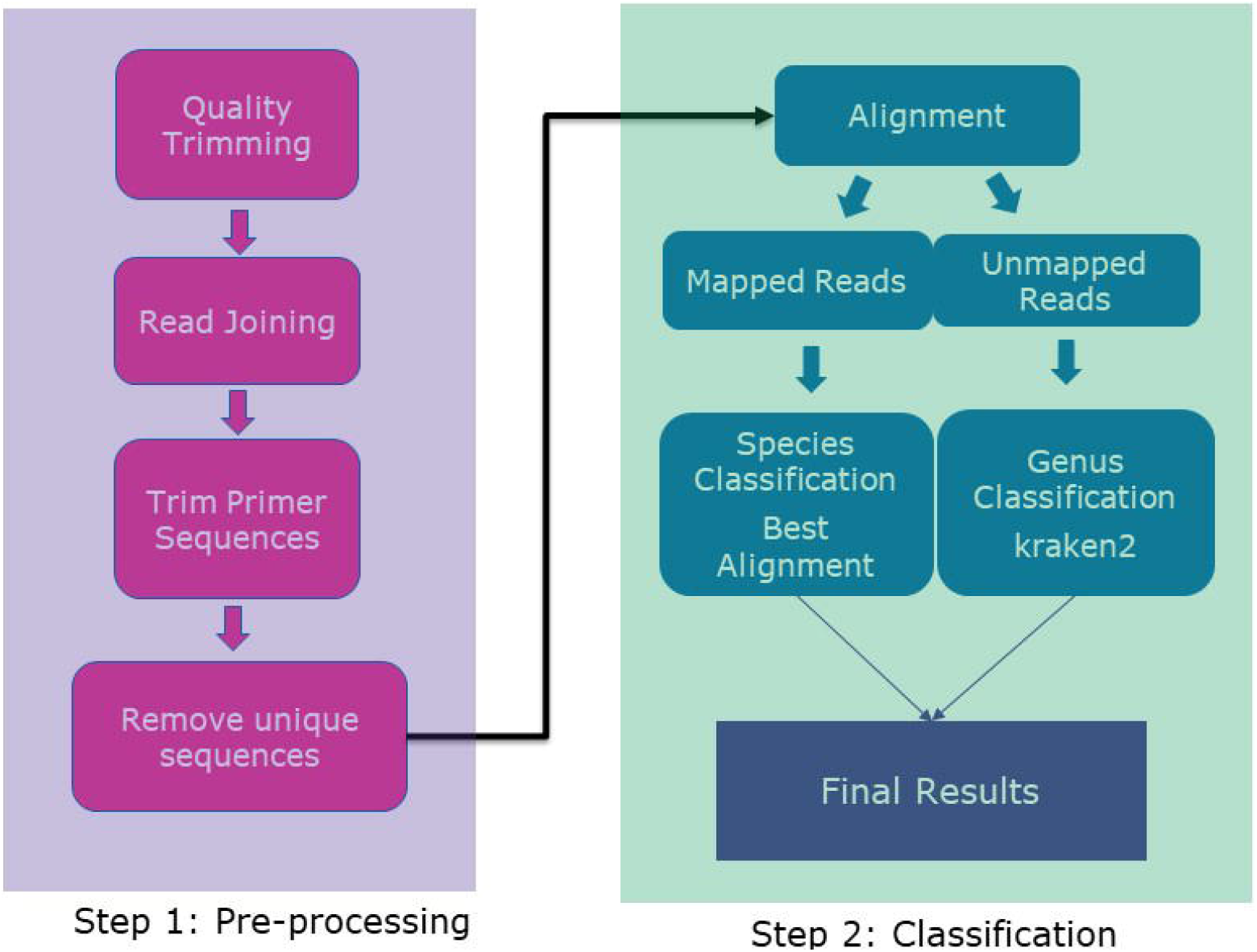
16S-seq Taxonomic Profiling Workflow. Workflow diagram of the 16S-seq taxonomic profiling software used in the M-CAMP Cloud Platform. The Pre-processing stage removes low quality reads and PCR artifacts which can lead to false positive classifications. The Classification stage uses alignments to the highly optimized M-CAMP 16S V3-V4 database to achieve high precision species level calls. A separate call to Kraken2 classifies reads that fail to align to the genus or family level. This second stage ensures that sequences from novel organisms are not left completely unclassified.

**Figure 2:**
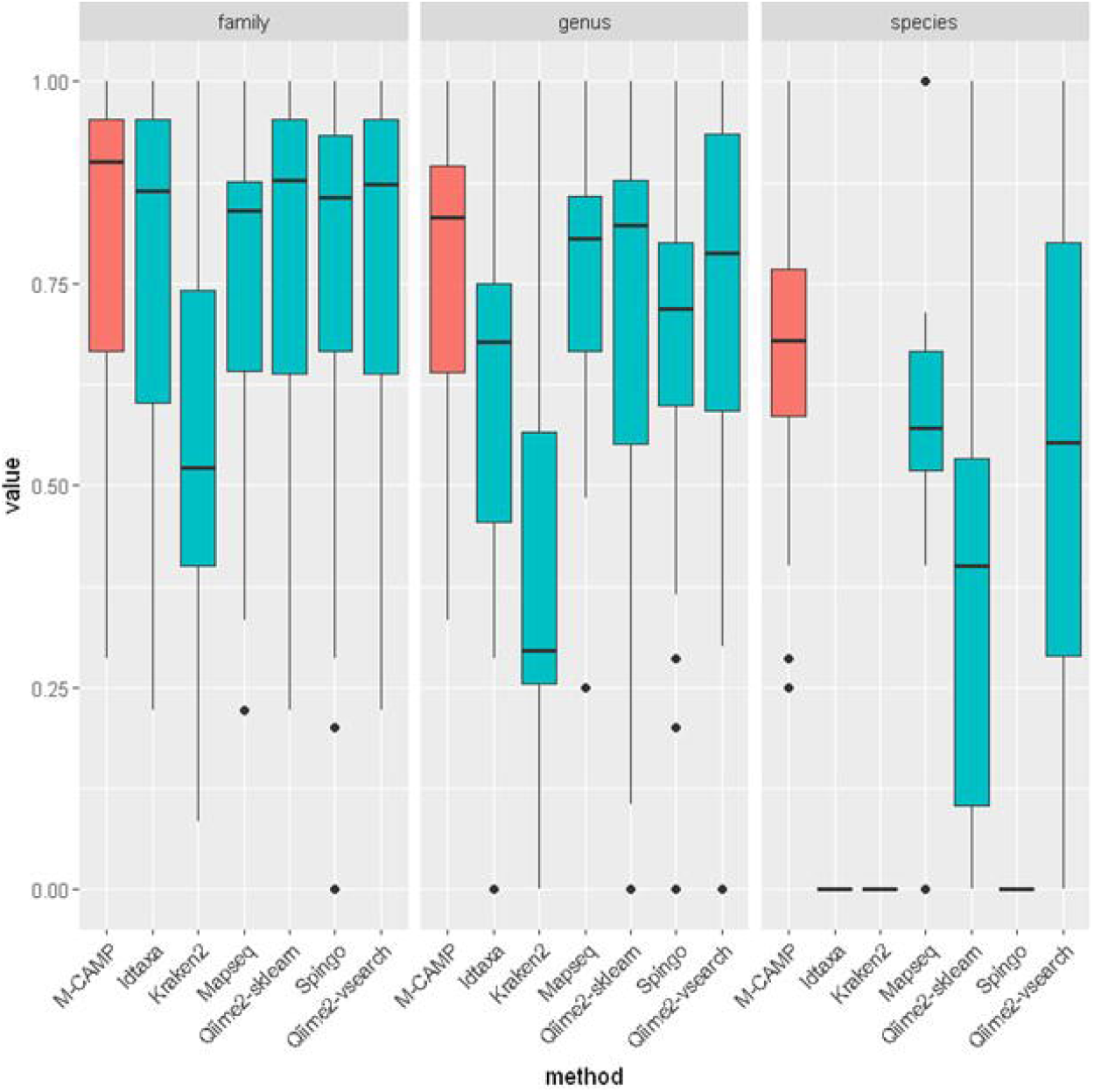
Boxplot of Fl Score. The Fl Score for different taxonomic levels were calculated for each of the validation samples and classification methods. The M-CAMP 16S method is shown in red. For all taxonomic levels, the median Fl score of the M-CAMP pipeline is the highest of all methods tested. For several of the tested methods the samples had a species-level Fl score of zero indicating the methods failed to make any species-level calls. This measure does not consider relative abundances

**Figure 3:**
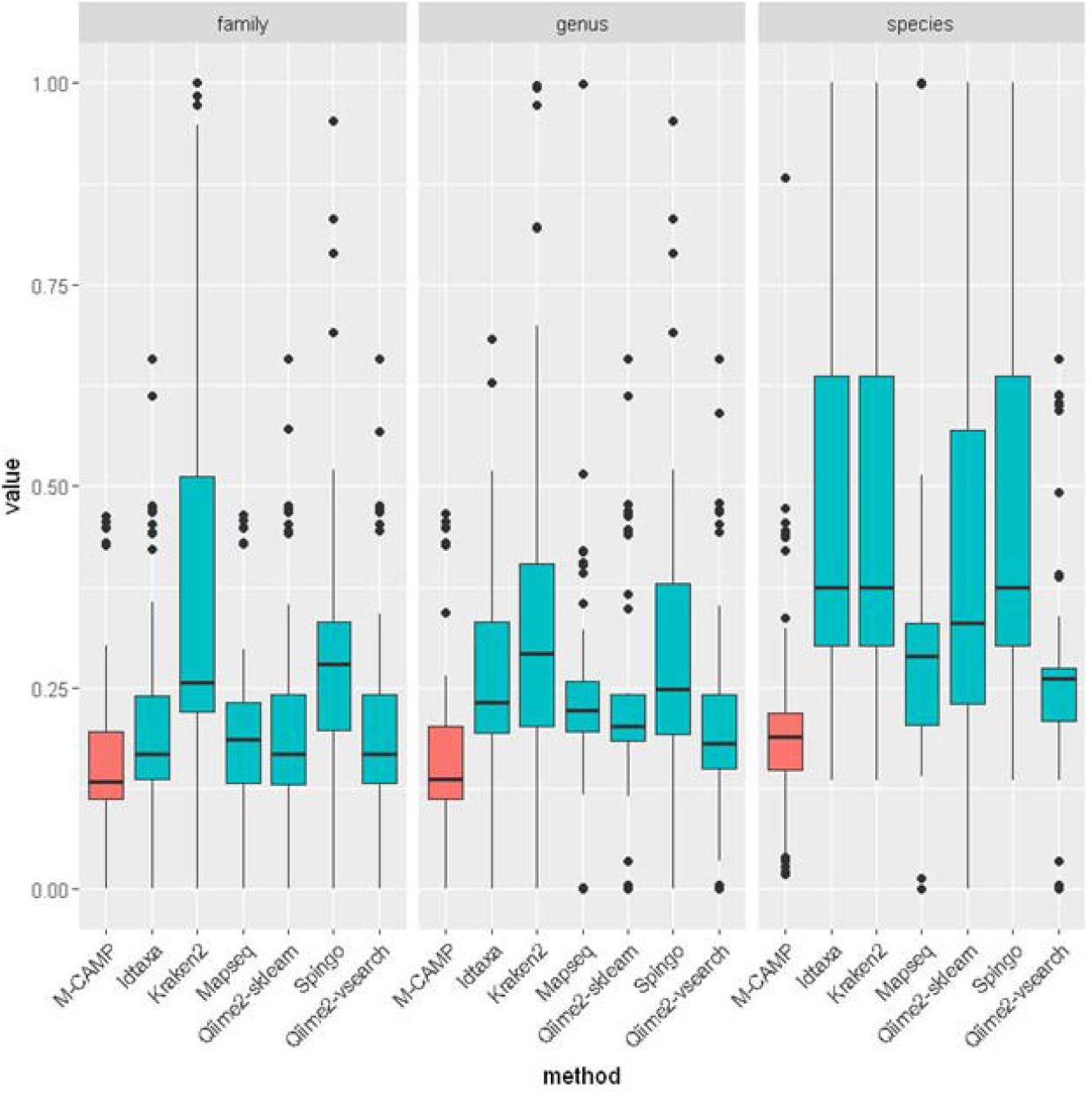
Boxplot of L2 Distance. Smaller values of the L2 distance indicate that the predicted and known taxonomic profiles are very similar. For each sample and classification method the L2 distance between the predicted and known bacterial relative abundances were calculated. The L2 distance is also known as the Euclidean distance. At all taxonomic levels, the M-CAMP 16S classifier has the best performance.

**Figure 4:**
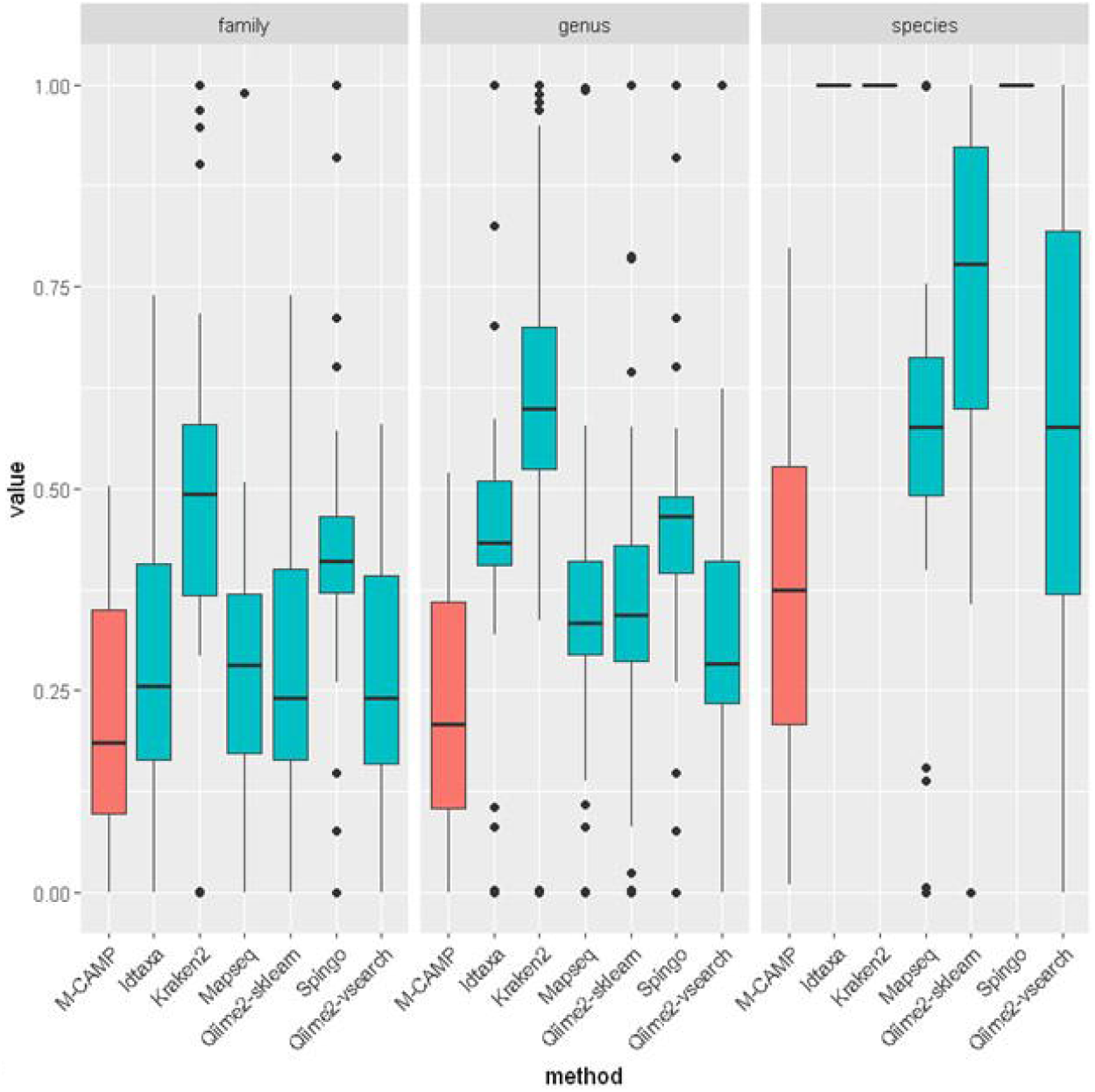
Boxplot of Bray-Curtis Distance. Lower Bray-Curtis distance is better. For each sample and classification method the Bray-Curtis distance between the predicted and known bacterial relative abundances were calculated. The Bray-Curtis distance is a popular non-phylogenetic measure of beta diversity. Smaller values of the distance mean that the predicted and known taxonomic profiles are very similar. For several of the tested methods the samples had a species-level distances of one indicating the methods failed to make any species-level calls. At all taxonomic levels, the M-CAMP16S classifier has the best performance.

**Figure 5:**
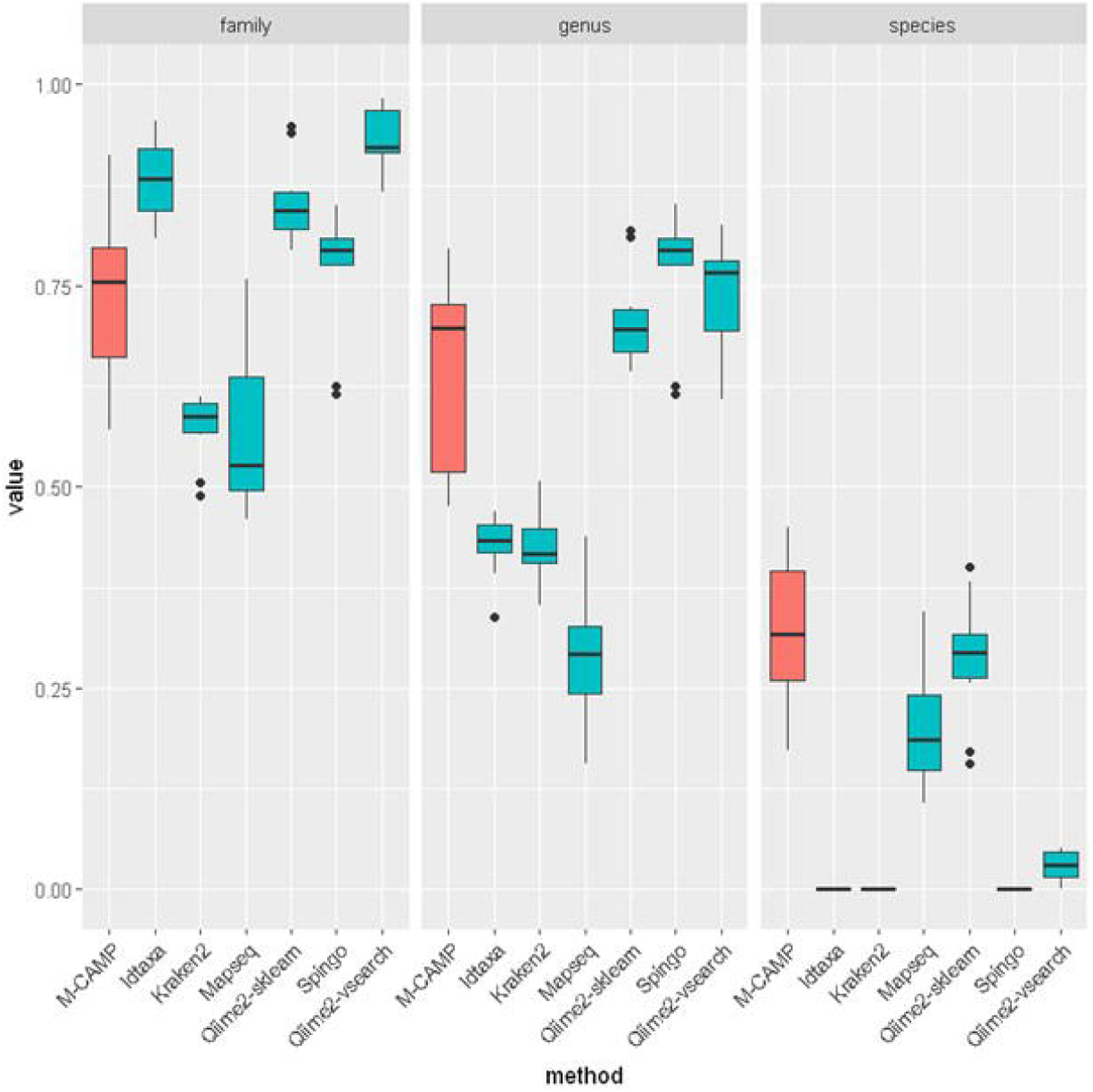
Percent of Reads Classified in Human Stool Samples. The fraction of reads from human stool samples that were classified different taxonomic levels. These stool samples contain an unknown microbial community so other performance metrics like Fl score cannot be calculated. On these complex real-world samples, the M-CAMP taxonomic classifier gives a high fraction of species and genus level classification.

The species level was where we saw the greatest level of improvement that the M-CAMP™ classification algorithm offers over the comparison methods. The M-CAMP™ 16s classification algorithm using the M-CAMP™ V3-V4 database has the lowest species level L2 distance of 0.213 compared to the next lowest Qiime2 vsearch with a L2 norm of 0.3055. M-CAMP™ 16s has the highest species level recall of 0.865 compared to the next highest of Mapseq run with the Mapref database with a recall of 0.526. M-CAMP™ 16s also has the highest species level F1 score of 0.865 compared to 0.526 for the next highest Mapref and 0.506 for Qiime2-vsearch. Three of the methods (Idtaxa, Kraken2, and Spingo) failed to give any species level calls and therefore had species level F1 scores of 0.

**Supplement Table S2** contains the known relative abundances for each validation sample. These known relative abundance values were used to calculate the evaluation metrics for the taxonomic profiles predicted by the classification methods.

### Statistical Comparison of Methods

We also performed a paired Wilcoxon test for the samples comparing all methods to the M-CAMP™ 16s classification algorithm at the species level. We focused the comparison at the species level because that is where we hope to provide the most improvement over existing methods. The shifts in the medians of each evaluation metric found by the test are shown in **Table 2**. All these shifts are relative to the M-CAMP™ 16s classification and they were all found to be statistically significant. For the L2 distance and Bray-Curtis distance a positive number represent decreased performance compared to the M-CAMP™ 16S classifier. For all other metrics a negative number represents decreased performance compared to the M-CAMP™ 16S classifier.

**Table 2:** Results from Paired Samples Wilcoxon Test. The value shown is the shift in the median of each metric compared to the M-CAMP™ 16S classifier. All shifts were found to be statistically significant at p<0.05. Positive values for L2 and Bray-Curtis represent decreased performance relative to the reference. Negative values for the other metrics represent decreased performance relative to the reference

| Metric | Idtaxa | Qiime2-vsearch | Qiime2-sklearn | Kraken2 | Spingo | Mapseq |
| --- | --- | --- | --- | --- | --- | --- |
| L2 | 0.149 | 0.06 | 0.114 | 0.149 | 0.149 | 0.063 |
| Bray-Curtis | 0.674 | 0.212 | 0.397 | 0.674 | 0.674 | 0.2 |
| Recall | -0.9 | -0.412 | -0.588 | -0.9 | -0.9 | -0.338 |
| Precision | -0.633 | 0.134 | -0.019 | -0.633 | -0.633 | 0.067 |
| F1 | -0.693 | -0.145 | -0.315 | -0.693 | -0.693 | -0.113 |

We found that the M-CAMP™ 16s classification algorithm outperforms all other algorithms in relative abundance estimation as measured by the L2 and Bray-Curtis distance, the ability to detect known bacteria as measured by the recall, and the overall accuracy of the taxonomic profile as measured by the F1 score. The Mapseq method has improved precision compared to M-CAMP™ algorithm, but with the cost of much lower recall. The L2 distance is the straight-line distance between the observed and true abundance vectors (Ye et al., 2019). We use the L2 distance because it is sensitive to the classification of high abundant taxa and it does not vary based on the phylogenetic distance. This allows us to evaluate the performance of our algorithm on microbiome community mixtures.

### Evaluation of Single Species Samples

Single species samples were included in our evaluation because they allow us to evaluate the performance of the classification methods on different phyla easily. **Table 3** shows the fraction of reads for each single bacteria sequencing sample that were classified to the correct bacteria at the species level. If a sample here has less than 100%, the rest of the reads were either misclassified or classified at the genus level or above. Only the M-CAMP™ 16s classification algorithm classifies above 90% of the reads to the correct species for most of the samples. For the M-CAMP™ 16s classification algorithm, the only sample that is below 90% is the Burkholderia pyrrocinia sample (Sigma Aldrich, Catalog MBD0019). However, none of the other methods were able to detect Burkholderia pyrrocinia, so the correct classification of at least 12% of the reads is a large improvement.

**Table 3:** Percent of Reads Correctly Classified at Species-level for Single Bacteria Samples.

| sample | species | M-CAMP 16S | Idtaxa | Qiime2-vsearch | Qiime2-sklearn | Kraken2 | Spingo | Mapseq |
| --- | --- | --- | --- | --- | --- | --- | --- | --- |
| BP2-1_S58 | Burkholderia pyrrocinia | 12.26% | 0% | 0.00% | 0.00% | 0.00% | 0.00% | 0% |
| BS1_S49 | Bacillus subtilis | 95.79% | 0% | 50.72% | 0.00% | 0.00% | 0.00% | 0% |
| EC_S7 | Escherichia coli | 94.50% | 0% | 99.95% | 0.00% | 0.00% | 0.00% | 0% |
| EF_S5 | Enterococcus faecalis | 97.86% | 0% | 99.98% | 0.00% | 0.00% | 0.00% | 0% |
| PA1_S52 | Pseudomonas aeruginosa | 94.43% | 0% | 99.51% | 0.00% | 0.00% | 0.00% | 0% |
| PG_S3 | Porphyromonas gingivalis | 97.25% | 0% | 99.98% | 99.98% | 0.00% | 0.00% | 100% |
| PM_S2 | Proteus mirabilis | 96.50% | 0% | 100.00% | 0.00% | 0.00% | 0.00% | 99% |
| PV_S1 | Proteus vulgaris | 96.06% | 0% | 0.00% | 0.00% | 0.00% | 0.00% | 0% |
| SE_S4 | Salmonella enterica | 98.14% | 0% | 97.55% | 0.00% | 0.00% | 0.00% | 76% |

The Qiime methods did not perform well in this test. The sklearn classifier did not achieve species level classification for any of the samples, and the vsearch classifier is only able to detect two species. Kraken2 performs well giving excellent results in four of the samples. Kraken2’s high species level recall comes at the cost of low species-level precision. As shown in Table 2, the average precision of the Kraken2 is lower than that achieved by the M-CAMP™ 16s classification algorithm. Mapseq detects just two of the species above the 90% read classification level.

Only the M-CAMP™ 16s classification algorithm delivered a species level classification for the E. coli sample. This is an expected weakness of most 16S-seq taxonomic classifiers because, as noted in the introduction, members of the Enterobacteriaceae family are known to have very similar 16S rRNA sequences (Grim et al., 2017). Our algorithm, along with the curated M-CAMP™ V3-V4 16S database, proved to be successful in the detection of species in this difficult to classify family.

### Evaluation of Runtime Performance

The runtime and resource usage of each method was recorded using the software GNU time version 1.7. **Figure 6** is a barplot showing the total user CPU time used for each of the algorithms. This time will be greater than walltime taken by the program if the program can run in multithreaded mode. When possible, we ran software with 32 threads per sample. Dada2 is a necessary pre-processing set for the Qiime2 and Idtaxa methods, which means the dada2 runtime must be added to the total runtime of those methods. Compared to the other methods, the total CPU time taken to run a sample through the M-CAMP™ 16s classification algorithm is very low, with only a couple of outlier samples which took longer to process. The performance of the algorithm is extremely important for the webapp service because the analysis must scale to include hundreds, or even thousands of samples run on commodity cloud servers.

**Figure 6:**
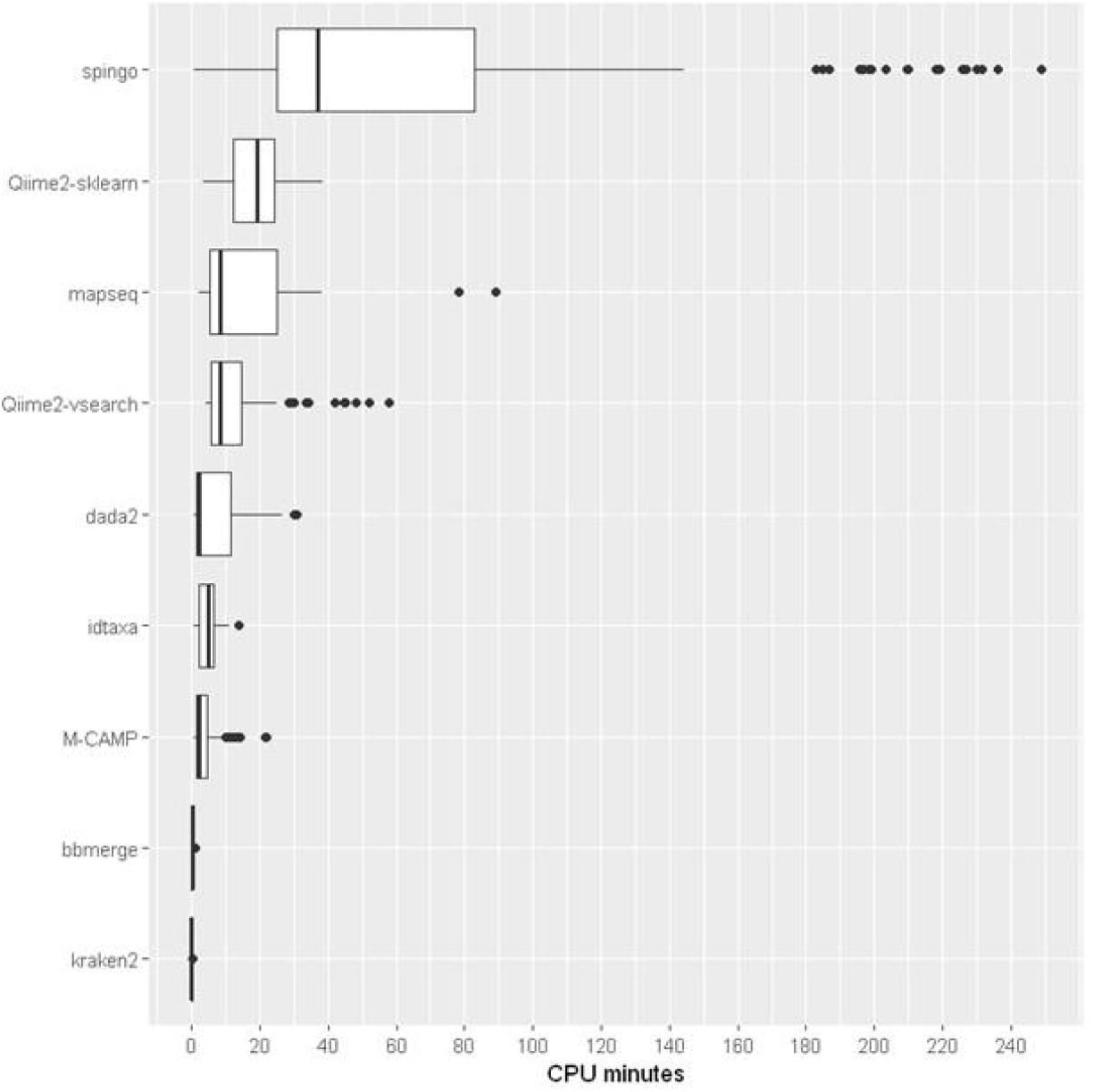
Runtimes of Software on Benchmark Samples. The number of CPU minutes needed for the classification software to process each validation sample. Preprocessing software like bbmerge and dada2 is included in this plot. CPU minutes is the walltime of the command multiplied by the number of CPUs utilized by the command. The M-CAMP 16S classifier is highly optimized and requires less time to run than most other methods.

## M-CAMP™ Cloud Platform

### Upload Fastq and Quality Assessment

The M-CAMP™ 16S taxonomic classification pipeline and database described here are available for public use on the M-CAMP™ platform. This platform provides registered users with intuitive graphical user interface to upload samples, run microbiome analysis, and view the results. **Figure 7** gives an overview of the functionality provided by the application. The sample management module in the web application allows users to create a new project with unique names, where fastq files for samples specific to the project can be uploaded. Additionally, users may opt to upload a metadata mapping file in tab delimited format describing experimental conditions of each sample in the project. The web platform facilitates users with options to perform taxonomic classification for 16S-seq bacterial amplicon sequencing, 18s-seq and ITS fungal amplicon sequencing, SMURF (Short MUltiple Regions Framework) (Fuks et al., 2018) for bacteria, and whole genome shotgun sequencing. The user can select the correct sequencing type during the fastq file upload process. To replicate the results of this paper, the user should select “Bacterial Amplicon-seq”.

**Figure 7:**
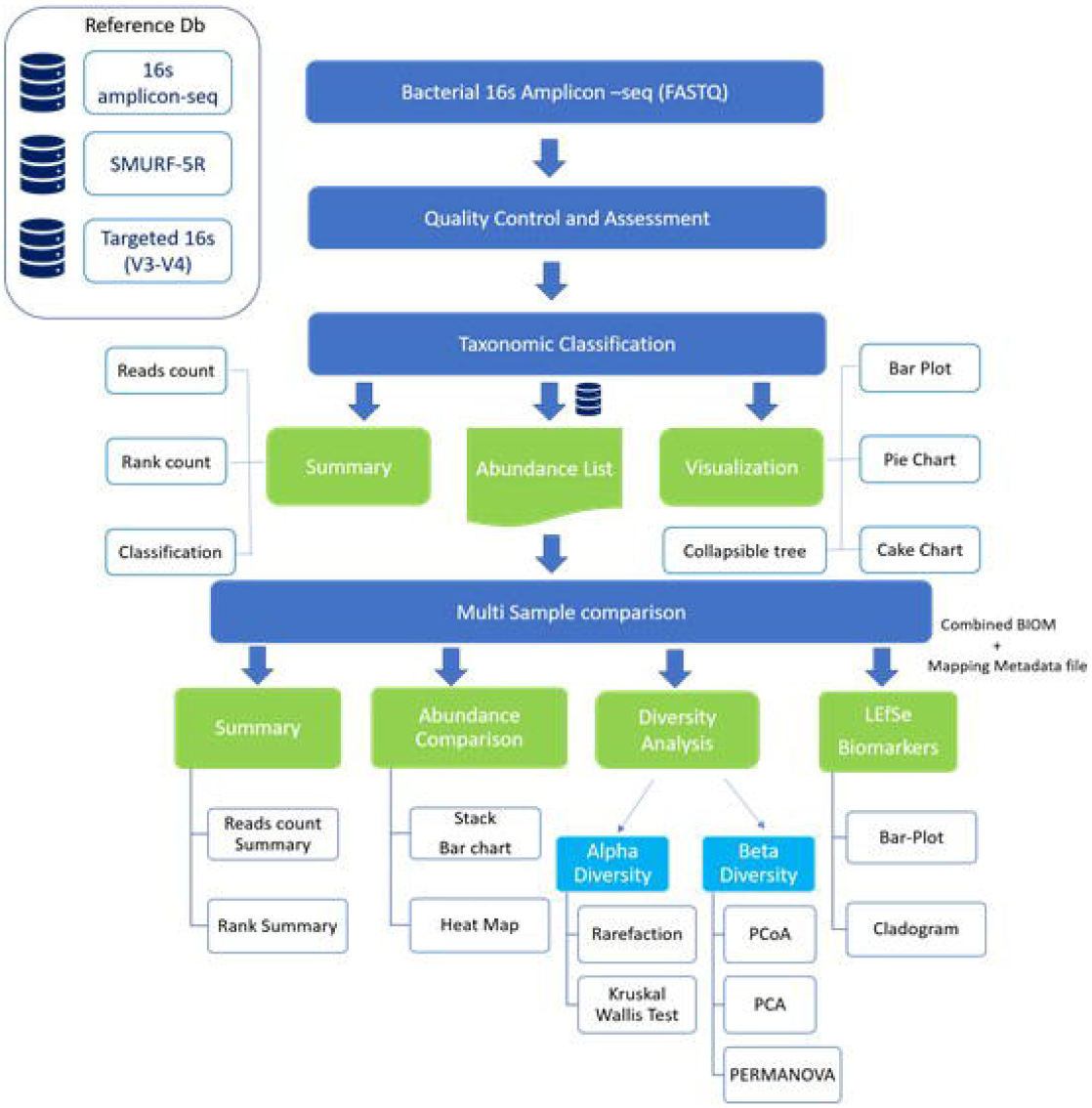
M-CAMP Cloud Platform Workflow. A list of modules and corresponding output files and graphics provided by the M-CAMP Cloud Platform. The Platform allows users to analyze single or paired-end Illumina generated fastq files

The uploaded fastq files undergo quality assessment automatically to generate preliminary reports of the sequencing quality for each of the uploaded samples. This quality report provides assessment on “per base sequence quality”, “per sequence quality score”, and “sequence length distribution”. The sample files are then processed using BBDuk to yield reads with average quality of Phred score(Q)>=30 (Bushnell et al., 2017). The user can also opt to set custom parameters for the trimming and filtering of all the fastq files in their project.

The post trimming and filtering quality assessment is auto performed by the application. The M-CAMP™ platform allows the user to view and download the QC report in the PDF or HTML format (**Figure 9A**) (TIBCO).

### Taxonomic Classification

The quality trimmed fastq files are used as inputs for taxonomic classification. At the classification step, users can select the reference database against which the classification needs to be performed. To use the database described in this paper, the user should select “M-CAMP-16S_V3-V4”. Once the classification status is “completed”, users can view the results online (**Figure 8**). Users can also download the sample classification report in the pdf format and/or in interactive HTML format (**Figure 9B**). The platform also allows users to download the required combined rank-based abundance file and other files consolidated in a zip format file at the sample level.

**Figure 8:**
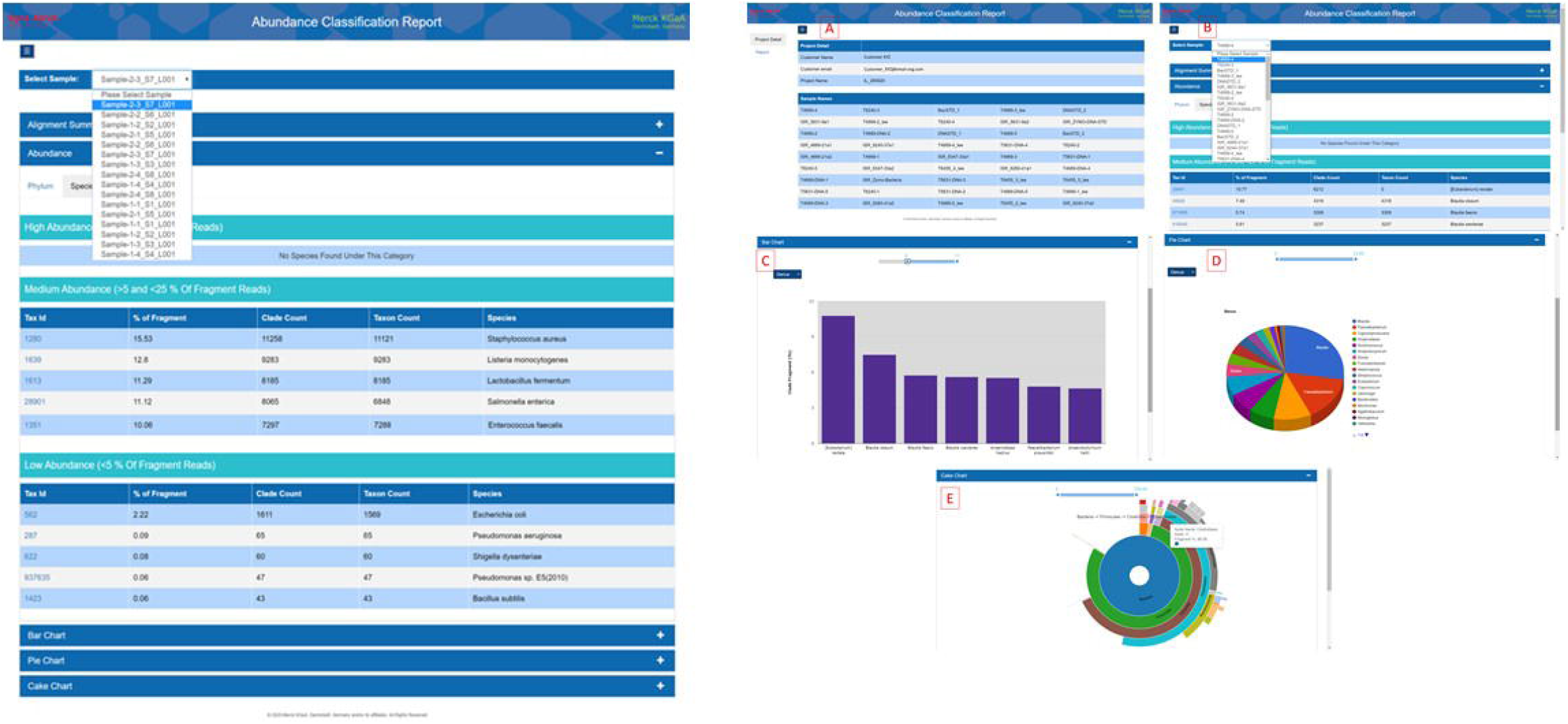
Downloadable HTML Reports. **A:** An example of downloadable HTML report for QC assessment **B** A example of downloadable HTML report for classification: A. Project level view; B. Sample level view; C-E. Sample level interactive graphics of classified fraction of bacteria

**Figure 9:**
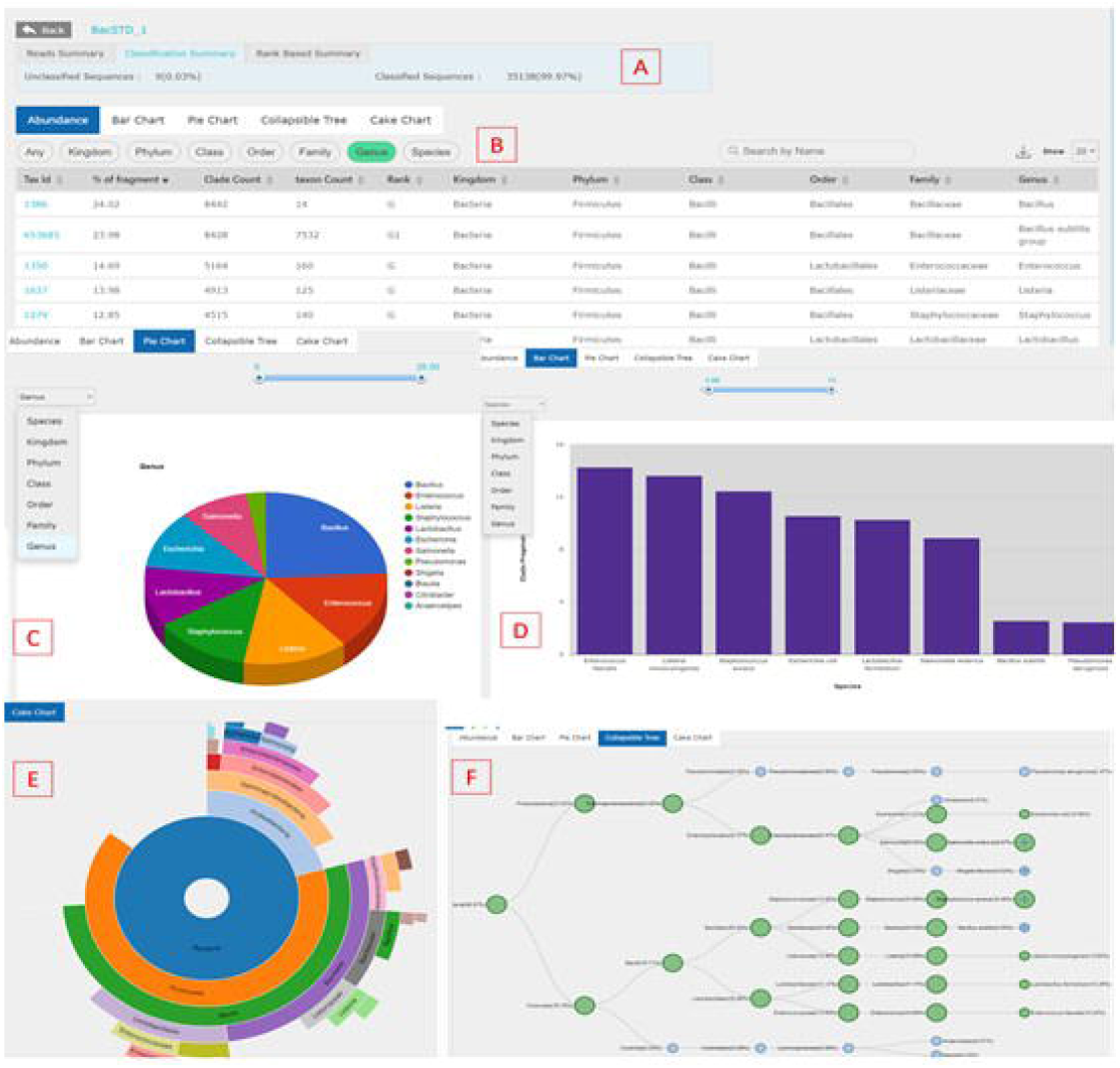
Taxonomic Classification Module Graphics. Taxonomic classification results and visualization. A. Reads counts, classification percentage and rank based summary B. Consolidated & rank wise relative abundance table C. Cut-off based interactive pie chart D. Rank-based interactive bar plot E. Interactive Cake Chart F. Abundance based collapsible hierarchical tree.

The platform allows users to share a project with any colleague that has an account on the platform. This allows users to collaboratively review the QC and taxonomic classification results. The collaborators with whom the project is shared can also download the raw fastq file of the shared project from the “Sample Management” module.

### Multiple sample comparative analysis

The comparative analysis module allows the user to perform diversity analysis using multiple samples within a selected project using information in the metadata table. Analysis in this module includes an overview of the classified read counts for each sample; alpha diversity, beta diversity, and barplots of the sample taxonomic profiles created using Qiime2 (Bolyen et al., 2019); heatmaps or abundant taxa in the project; and statistical tests for differentially abundant microbes using LEfSe (Segata et al., 2011). The multiple sample comparison analysis results can be viewed within the application or can be downloaded as a PDF or HTML report (**Figure 10**).

**Figure 10:**
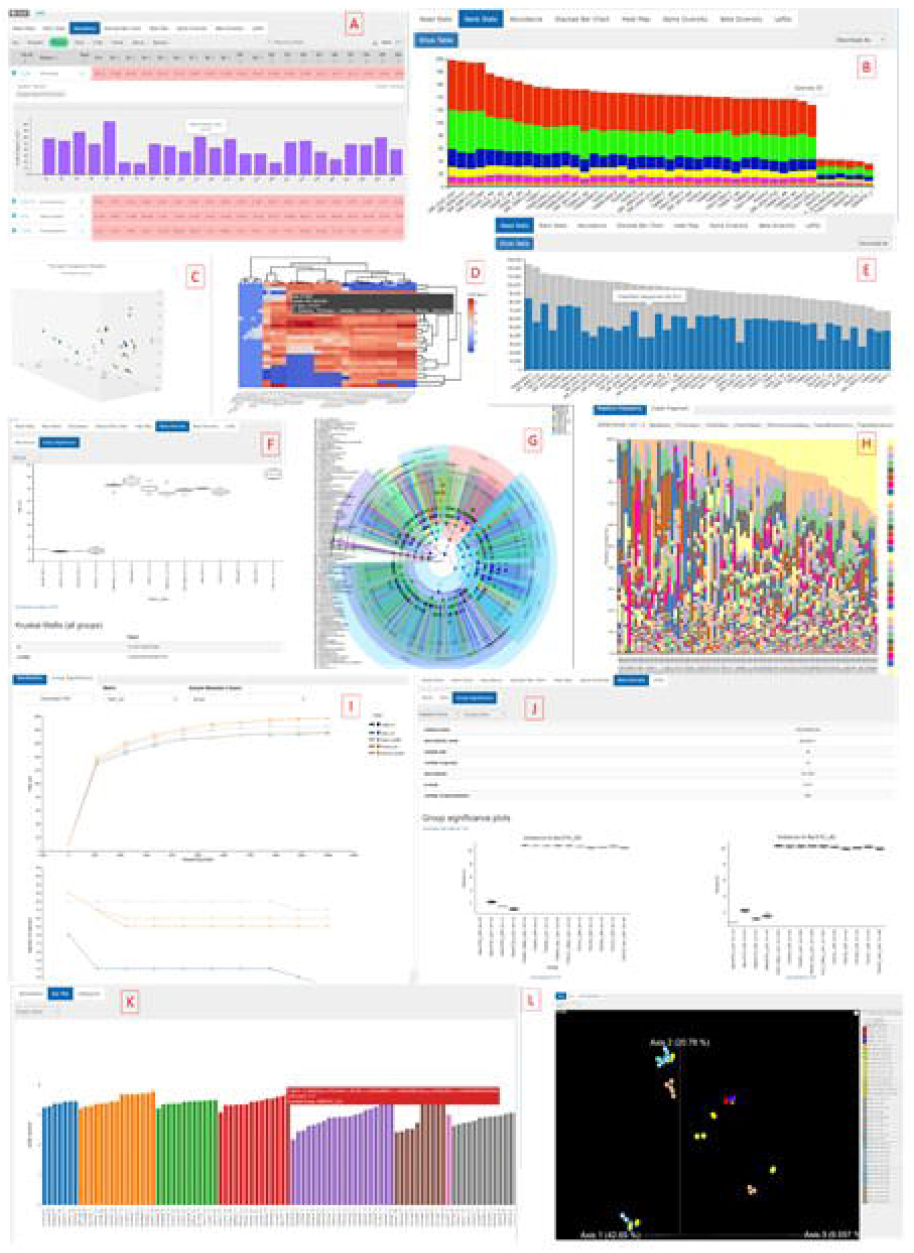
Comparative Analysis Module Graphics. A. Abundance (clade fragments count) percentage based multiple sample comparison table. B. Rank stats summarizing the count of taxa present in each sample at each rank level. C. PCA plot based upon the OTU frequency at the taxonomy rank level across samples D. Interactive heatmap based upon the log transformed CLR normalized abundance value at the species level. E. Classified and Un-classified reads count comparison across the sample F. Beta diversity Group significance using Permanova test for the specified distance metric G. Metadata class-based taxon abundance distribution H. Stacked bar-chart based upon the relative frequency I. Alpha rarefaction J. Kruskal-Wallistest based Alpha group significance K. Bar plots showing LEfSe based class differences across samples. L. PCoA for the multivariate analysis for Beta-diversity. PcoA visualization for the categorized samples can be seen based upon the Bray-curtis, Jaccard, Weighted Unifrac and Un-weighted Unifrac metrics respectively.

### Evaluation of Sigma-Aldrich Microbial Community DNA Mix

The Standard Mix module allows users to evaluate the accuracy of taxonomic classification on positive control community mixtures. Running positive control samples is important to verify that no errors are introduced by the DNA extraction or sequencing methods. Positive samples are also crucial for assessing the accuracy of bioinformatics methods for taxonomic classification. With so many variables in the experimental and analytical processing of a microbiome project, these control samples are crucial for ensuring that the results are reproducible (Hornung, Zwittink, & Kuijper, 2019).

The Sigma-Aldrich Microbial community DNA mix allows researchers to include high-quality positive controls in their projects without having to create their own synthetic community mixtures. (Microbial community DNA mix, Sigma-Aldrich, Catalog: MBD0026-0.3UG). This community contains genomic DNA from 10 different bacterial species with different genome sizes, GC content, and phyla. **Figure 11** shows the report generated by the Standard Mix module of the M-CAMP™ Cloud Platform on one of the Sigma-Aldrich microbiome samples named in this study as STD-D-sigma. The metrics in this online report are the same as those presented in this manuscript. The values are calculated for each of the seven main taxonomic levels. At the species level the M-CAMP™ 16s classification algorithm has almost perfect F1 score and low Bray-Curtis distance. **Figure 12** shows another report detailing the specific taxonomic classifications with relative abundances for the sample. This report shows that all 10 species were successfully detected by the Cloud Platform taxonomic classification pipeline with two low abundance false positive species detected.

**Figure 11:**
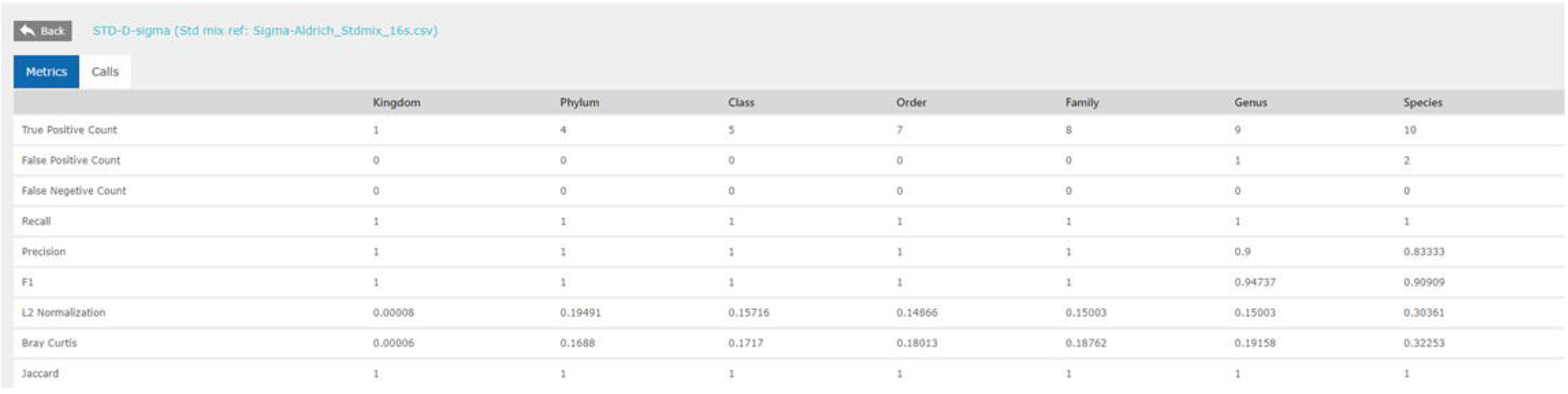
Standards Mix Module Report. An example report generated by the Standard Mix module of the M-CAMP Cloud Platform. The sample being evaluated is sequencing of the Sigma-Aldrich Microbial Community DNA mix. This report shows performance metrics calculated at the seven main taxonomic levels.

**Figure 12:**
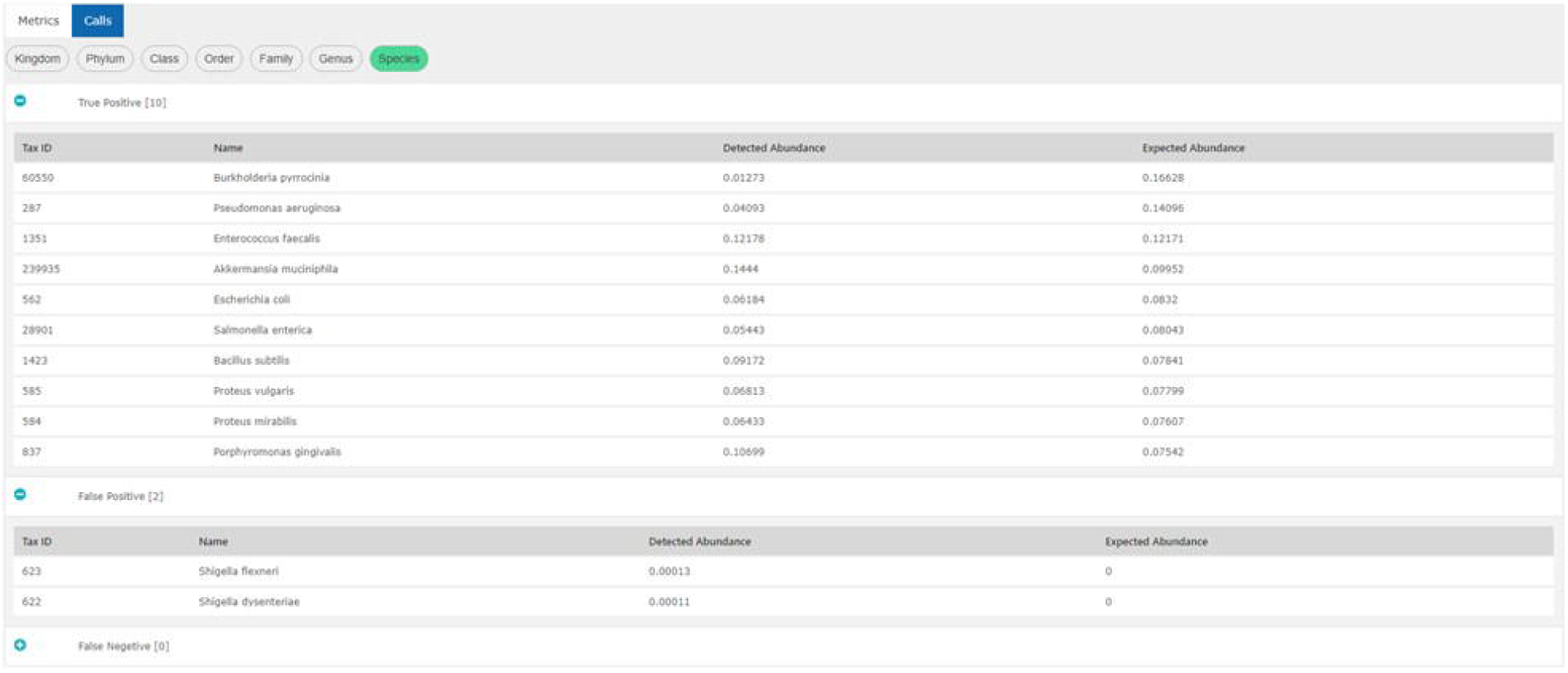
Standards Mix Module Taxonomic Report. An example report generated by the Standard Mix module for a Sigma-Aldrich Microbial Community DNA Mix sample. This report shows the taxonomic calls made by the Classification module and groups them by “True Positive”, “False Positive” and “False Negative”. Predicted and expected relative abundances are show for each taxonomy. The taxonomic level can be selected by clicking the bubbles at the top.

## Discussion

We have presented a novel 16s taxonomic classification pipeline that is unique because it combines the two most common approaches to 16S classification: kmer classification and alignment to reference sequences. This combination allows the strengths of both approaches to cover each other’s weaknesses. We have also presented a novel method of combined multiple 16S sequence databases and a method of curating the associated taxonomic labels. We have shown that the M-CAMP™ 16S taxonomic classification algorithm run with a curated V3V4 region database is able to classify 16S-seq Illumina data to the species level. This method consistently outperforms state of the art methods on synthetic microbiome community samples.

The taxonomic classification pipeline is available to the public on the M-CAMP™ Cloud Platform. The cloud platform allows researchers to upload their metagenome sequencing data and generate publication quality graphs and statistical analysis without any having the bioinformatics skills or computer hardware normally required to run such projects. The interactive reports generated by the platform can be view online in the platform or downloaded for offline viewing. Access to the platform can be requested here: https://www.sigmaaldrich.com/life-science/microbiome.html.

## Availability of data and materials

The sequencing data used for this study along with the supplemental tables and methods are available here https://doi.org/10.17632/xyhphwr4b2.1

## Conflict of Interest

Authors are employed by MilliporeSigma which develops the M-CAMP™ Cloud Platform.

The authors acknowledge the important contributions of of Graziella Amarasinghe, Liron Beladev, Anjani Kumar Deepak, and Yi Sun in making this manuscript possible.

## References

Allard, G., Ryan, F. J., Jeffery, I. B., & Claesson, M. J. (2015). SPINGO: a rapid species-classifier for microbial amplicon sequences. BMC Bioinformatics, 16, 324. doi:10.1186/s12859-015-0747-1

Bolyen, E., Rideout, J. R., Dillon, M. R., Bokulich, N. A., Abnet, C. C., Al-Ghalith, G. A., … Caporaso, J. G. (2019). Reproducible, interactive, scalable and extensible microbiome data science using QIIME 2. Nat Biotechnol, 37(8), 852–857. doi:10.1038/s41587-019-0209-9

Boyer, F., Mercier, C., Bonin, A., Le Bras, Y., Taberlet, P., & Coissac, E. (2016). obitools: a unix-inspired software package for DNA metabarcoding. Mol Ecol Resour, 16(1), 176–182. doi:10.1111/1755-0998.12428

Bushnell, B., Rood, J., & Singer, E. (2017). BBMerge - Accurate paired shotgun read merging via overlap. PLoS One, 12(10), e0185056. doi:10.1371/journal.pone.0185056

Caporaso, J. G., Kuczynski, J., Stombaugh, J., Bittinger, K., Bushman, F. D., Costello, E. K., … Knight, R. (2010). QIIME allows analysis of high-throughput community sequencing data. Nat Methods, 7(5), 335–336. doi:10.1038/nmeth.f.303

Cole, J. R., Wang, Q., Fish, J. A., Chai, B., McGarrell, D. M., Sun, Y., … Tiedje, J. M. (2014). Ribosomal Database Project: data and tools for high throughput rRNA analysis. Nucleic Acids Res, 42(Database issue), D633–642. doi:10.1093/nar/gkt1244

Council, N. R. (2007). In The New Science of Metagenomics: Revealing the Secrets of Our Microbial Planet. Washington (DC).

DeSantis, T. Z., Hugenholtz, P., Larsen, N., Rojas, M., Brodie, E. L., Keller, K., … Andersen, G. L. (2006). Greengenes, a chimera-checked 16S rRNA gene database and workbench compatible with ARB. Appl Environ Microbiol, 72(7), 5069–5072. doi:10.1128/AEM.03006-05

Fettweis, J. M., Serrano, M. G., Brooks, J. P., Edwards, D. J., Girerd, P. H., Parikh, H. I., … Buck, G. A. (2019). The vaginal microbiome and preterm birth. Nat Med, 25(6), 1012–1021. doi:10.1038/s41591-019-0450-2

Fuks, G., Elgart, M., Amir, A., Zeisel, A., Turnbaugh, P. J., Soen, Y., & Shental, N. (2018). Combining 16S rRNA gene variable regions enables high-resolution microbial community profiling. Microbiome, 6(1), 17. doi:10.1186/s40168-017-0396-x

Gohl, D. M., Vangay, P., Garbe, J., MacLean, A., Hauge, A., Becker, A., … Beckman, K. B. (2016). Systematic improvement of amplicon marker gene methods for increased accuracy in microbiome studies. Nat Biotechnol, 34(9), 942–949. doi:10.1038/nbt.3601

Grim, C. J., Daquigan, N., Lusk Pfefer, T. S., Ottesen, A. R., White, J. R., & Jarvis, K. G. (2017). High-Resolution Microbiome Profiling for Detection and Tracking of Salmonella enterica. Front Microbiol, 8, 1587. doi:10.3389/fmicb.2017.01587

Gruning, B., Dale, R., Sjodin, A., Chapman, B. A., Rowe, J., Tomkins-Tinch, C. H., … Bioconda, T. (2018). Bioconda: sustainable and comprehensive software distribution for the life sciences. Nat Methods, 15(7), 475–476. doi:10.1038/s41592-018-0046-7

Hanson, N. W., Konwar, K. M., & Hallam, S. J. (2016). LCA*: an entropy-based measure for taxonomic assignment within assembled metagenomes. Bioinformatics, 32(23), 3535–3542. doi:10.1093/bioinformatics/btw400

Herlemann, D. P., Labrenz, M., Jurgens, K., Bertilsson, S., Waniek, J. J., & Andersson, A. F. (2011). Transitions in bacterial communities along the 2000 km salinity gradient of the Baltic Sea. ISME J, 5(10), 1571–1579. doi:10.1038/ismej.2011.41

Hornung, B. V. H., Zwittink, R. D., & Kuijper, E. J. (2019). Issues and current standards of controls in microbiome research. FEMS Microbiol Ecol, 95(5). doi:10.1093/femsec/fiz045

Huerta-Cepas, J., Serra, F., & Bork, P. (2016). ETE 3: Reconstruction, Analysis, and Visualization of Phylogenomic Data. Mol Biol Evol, 33(6), 1635–1638. doi:10.1093/molbev/msw046

Lloyd-Price, J., Arze, C., Ananthakrishnan, A. N., Schirmer, M., Avila-Pacheco, J., Poon, T. W., … Huttenhower, C. (2019). Multi-omics of the gut microbial ecosystem in inflammatory bowel diseases. Nature, 569(7758), 655–662. doi:10.1038/s41586-019-1237-9

Matias Rodrigues, J. F., Schmidt, T. S. B., Tackmann, J., & von Mering, C. (2017). MAPseq: highly efficient k-mer search with confidence estimates, for rRNA sequence analysis. Bioinformatics, 33(23), 3808–3810. doi:10.1093/bioinformatics/btx517

Murali, A., Bhargava, A., & Wright, E. S. (2018). IDTAXA: a novel approach for accurate taxonomic classification of microbiome sequences. Microbiome, 6(1), 140. doi:10.1186/s40168-018-0521-5

O’Leary, N. A., Wright, M. W., Brister, J. R., Ciufo, S., Haddad, D., McVeigh, R., … Pruitt, K. D. (2016). Reference sequence (RefSeq) database at NCBI: current status, taxonomic expansion, and functional annotation. Nucleic Acids Res, 44(D1), D733–745. doi:10.1093/nar/gkv1189

Quast, C., Pruesse, E., Yilmaz, P., Gerken, J., Schweer, T., Yarza, P., … Glockner, F. O. (2013). The SILVA ribosomal RNA gene database project: improved data processing and web-based tools. Nucleic Acids Res, 41(Database issue), D590–596. doi:10.1093/nar/gks1219

Rognes, T., Flouri, T., Nichols, B., Quince, C., & Mahé, F. (2016). VSEARCH: a versatile open source tool for metagenomics. PeerJ, 4, e2584. doi:10.7717/peerj.2584

Schirmer, M., Ijaz, U. Z., D’Amore, R., Hall, N., Sloan, W. T., & Quince, C. (2015). Insight into biases and sequencing errors for amplicon sequencing with the Illumina MiSeq platform. Nucleic Acids Res, 43(6), e37. doi:10.1093/nar/gku1341

Seemann, T. (2013). Barrnap (Version 0.9). Retrieved from https://github.com/tseemann/barrnap

Segata, N., Izard, J., Waldron, L., Gevers, D., Miropolsky, L., Garrett, W. S., & Huttenhower, C. (2011). Metagenomic biomarker discovery and explanation. Genome Biology, 12(6), 1-18. doi:doi:10.1186/gb-2011-12-6-r60

TIBCO. JasperReports: TIBCO Software Inc.

Tourlousse, D. M., Yoshiike, S., Ohashi, A., Matsukura, S., Noda, N., & Sekiguchi, Y. (2017). Synthetic spike-in standards for high-throughput 16S rRNA gene amplicon sequencing. Nucleic Acids Res, 45(4), e23. doi:10.1093/nar/gkw984

Tyson, G. W., Chapman, J., Hugenholtz, P., Allen, E. E., Ram, R. J., Richardson, P. M., … Banfield, J. F. (2004). Community structure and metabolism through reconstruction of microbial genomes from the environment. Nature, 428(6978), 37–43. doi:10.1038/nature02340

Venter, J. C., Remington, K., Heidelberg, J. F., Halpern, A. L., Rusch, D., Eisen, J. A., … Smith, H. O. (2004). Environmental genome shotgun sequencing of the Sargasso Sea. Science, 304(5667), 66–74. doi:10.1126/science.1093857

Woese, C. R. (1987). Bacterial evolution. Microbiol Rev, 51(2), 221–271.

Wood, D. E., Lu, J., & Langmead, B. (2019). Improved metagenomic analysis with Kraken 2. Genome Biol, 20(1), 257. doi:10.1186/s13059-019-1891-0

Wood, D. E., & Salzberg, S. L. (2014). Kraken: ultrafast metagenomic sequence classification using exact alignments. Genome Biol, 15(3), R46. doi:10.1186/gb-2014-15-3-r46

Ye, S. H., Siddle, K. J., Park, D. J., & Sabeti, P. C. (2019). Benchmarking Metagenomics Tools for Taxonomic Classification. Cell, 178(4), 779–794. doi:10.1016/j.cell.2019.07.010

Zhang, X., Shao, Y., Tian, J., Liao, Y., Li, P., Zhang, Y., … Li, Z. (2019). pTrimmer: An efficient tool to trim primers of multiplex deep sequencing data. BMC Bioinformatics, 20(1), 236. doi:10.1186/s12859-019-2854-x

Zhou, W., Sailani, M. R., Contrepois, K., Zhou, Y., Ahadi, S., Leopold, S. R., … Snyder, M. (2019). Longitudinal multi-omics of host-microbe dynamics in prediabetes. Nature, 569(7758), 663–671. doi:10.1038/s41586-019-1236-x

Gao, B., Chi, L., Zhu, Y., Shi, X., Tu, P., Li, B., … Schnabl, B. (2021). An Introduction to Next Generation Sequencing Bioinformatic Analysis in Gut Microbiome Studies. Biomolecules, 11(4). doi:10.3390/biom11040530

Nearing, J. T., Comeau, A. M., & Langille, M. G. I. (2021). Identifying biases and their potential solutions in human microbiome studies. Microbiome, 9(1), 113. doi:10.1186/s40168-021-01059-0

