## Supplemental Methods for "M-CAMP™: A cloud-based web platform with a novel approach for species-level classification of 16S rRNA microbiome sequences"

**Detailed Benchmark Commands**

Qiime2 [9] was run using version 2019.10. Fastq files were processed with dada2 using this command:

qiime dada2 denoise-paired --p-n-threads 1 \

--i-demultiplexed-seqs $tmpdir/demux.qza \

--p-trim-left-f 23 \

--p-trim-left-r 23 \

--p-trunc-len-f <read_length> \

--p-trunc-len-r <read_length> \

--p-chimera-method pooled \

--o-representative-sequences seqs.qza \

--o-table table.qza \

--o-denoising-stats $tmpdir/dada2-stats.qza

Both the sklearn and vsearch classifier available in Qiime2 were tested using the Silva 138 database as the reference. Three sklearn models were trained using different sets of primers representing different amplicons. For each sample in the benchmarking dataset the model trained with the primers used in the 16S-seq protocol were used. The primer specific representative sequences for the appropriate amplicon were also use for the vsearch classifier.

CCTACGGGNGGCWGCAG GACTACHVGGGTATCTAATCC V3V4

GTGYCAGCMGCCGCGGTAA GGACTACNVGGGTWTCTAAT V4

Training was performed with the following command:

qiime feature-classifier extract-reads \

--i-sequences rep_set.qza \

--p-f-primer $FWD_PRIMER \

--p-r-primer $REV_PRIMER \

--o-reads $OUTDIR/ref-seqs.qza

qiime feature-classifier fit-classifier-naive-bayes \

--i-reference-reads $OUTDIR/ref-seqs.qza \

--i-reference-taxonomy ref-taxonomy.qza \

--o-classifier $OUTDIR/nb-classifier.qza

The sklearn classifier was run with this command:

qiime feature-classifier classify-sklearn \

--p-n-jobs <threads> \

--i-classifier <clf_dir>/nb-classifier.qza \

--i-reads seqs.qza \

--o-classification <output_taxa>\

--p-confidence 0.7

The vsearch classifier was run with reference sequences from the Silva 138 database with this command:

qiime feature-classifier classify-consensus-vsearch \

--i-query seqs.qza \

--i-reference-reads <database_dir>/rep_set.qza \

--i-reference-taxonomy <database_dir>/ref-taxonomy.qza \

--p-threads <self.threads> \

--o-classification <output_taxa>

Spingo was installed from the bioconda package spingo==1.3=py27_boost1.64. Spingo was run with default database RDP_11.2.species.fa described here <https://github.com/GuyAllard/SPINGO/tree/master/database>. Input sequences were quality trimmed and had primer sequences removed with trimmomatic and pair merged using bbmerge. Classification was performed with the following command:

spingo --write-index -d <database> -i <data[0]> -p <threads> > <read_clf>

spingo_summary.py <read_clf> --level 2 > <report>

Mapseq version mapseq-1.2.3-linux was installed from the precompiled binary on the author’s website here <https://www.meringlab.org/software/mapseq/> . The database mapref-2.2b was used in the classification. Input sequences were quality trimmed and had primer sequences removed with trimmomatic and pair merged using bbmerge. Classification was run using this command:

mapseq --nthreads <threads> --outfmt simple <data> > <mseq>

Idtaxa was installed from bioconda package bioconductor-decipher==2.16.0=r40h037d062_0. Input reads were processed by the qiime2 dada2 denoise-paired command. The database used was SILVA_SSU_r138_2019 downloaded from the DECIPHER project page here <http://www2.decipher.codes/Downloads.html>. The Rscript used to classify is named idtaxa.R (link to location)

Kraken2 was installed from bioconda package kraken2== 2.0.8_beta=pl526h6bb024c_1. Input reads were quality trimmed and had primer sequences removed with trimmomatic and pair merged using bbmerge. Classification was run using this command:

kraken2 --db <silva database> --threads 8 --report <output report> --confidence 0.2
